# OTX2 non-cell autonomous activity regulates inner retinal function

**DOI:** 10.1101/2020.02.12.945584

**Authors:** Raoul Torero-Ibad, Bilal Mazhar, Clémentine Vincent, Clémence Bernard, Julie Dégardin, Manuel Simonutti, Thomas Lamonerie, Ariel Di Nardo, Alain Prochiantz, Kenneth L. Moya

**Author notes:** Correspondence should be addressed to: Kenneth L. Moya. These authors contributed equally to this work (Raoul Torero-Ibad, Bilal Mazhar, Clémentine Vincent). Author contributions: RTI, BM, CV, CB, JD, MS performed research; RTI, CV, CB, ADN and KLM analyzed data; RTI, TL, ADN, AP and KLM designed research; RTI, AP and KLM wrote the paper.

## Abstract

OTX2 is a homeoprotein transcription factor expressed in photoreceptors and bipolar cells in the retina. OTX2, like many other homeoproteins, transfers between cells and exerts non-cell autonomous effects such as promoting survival of retinal ganglion cells that do not express the protein. Here we used a genetic approach to target extracellular OTX2 in the retina by conditional expression of a secreted single chain anti-OTX2 antibody. Compared to control mice, the expression of this antibody by Parvalbumin-expressing neurons in the retina is followed by a reduction in visual acuity in one-month-old mice with no alteration of the retinal structure or cell type number or aspect. A- and b-waves measured by electroretinogram were also indistinguishable from control mice, suggesting no functional deficit of photoreceptors and bipolar cells. Mice expressing the OTX2-neutralizing antibody did show a significant doubling in the flicker amplitude, consistent with a change in inner retinal function. Our results show that interfering *in vivo* with OTX2 non-cell autonomous activity in the postnatal retina leads to an alteration in inner retinal cell functions and causes a deficit in visual acuity.

**Significance statement:** OTX2 is a homeoprotein transcription factor expressed in retinal photoreceptors and bipolar cells. Although the *Otx2* locus is silent in the inner retina, the protein is detected in cells of the ganglion cell layer consistent with the ability of this class of proteins to transfer between cells. We expressed a secreted single chain antibody (scFv) against OTX2 in the retina to neutralize extracellular OTX2. Antibody expression leads to reduced visual acuity with no change in retinal structure, or photoreceptor or bipolar physiology; however, activity in the inner retina was altered. Thus, interfering with OTX2 non-cell autonomous activity in postnatal retina alters inner retinal function and causes vision loss, highlighting the physiological value of homeoprotein direct non-cell autonomous signaling.

## Introduction

OTX2 is a homeoprotein transcription factor important for retinal development and maintenance. It is expressed early in the embryonic mouse optic vesicle and retinal pigmented epithelium (RPE) and is required for differentiation of photoreceptors by transactivation of *Crx* and *Nrl* and for differentiation of bipolar cells (BPs) via regulation of PKCα (Martinez-Morales et al, 2001; Nishida et al, 2003; Koike, et al. 2007; Murinashi et al., 2011). In mice hypomorph for *Otx2* we reported visual deficits, retinal physiological dysfunction and age- and OTX2-dependent degeneration of the retina (Bernard et al., 2014). In adult mice, knock down of *Otx2* in RPE results in photoreceptor death demonstrating that continued expression of OTX2 in RPE is necessary for photoreceptor survival (Housset et al., 2013). Exogenous OTX2 protects adult retinal galnglion cells (RGCs) against NMDA-induced excitotoxicity and preserves visual acuity (Torero Ibad et al., 2011).

The capacity of homeoproteins (HPs) to transfer between cells allows different types of activities. Homeoproteins can act within the cells that produce them, thus in a cell autonomous fashion, but they can also exert their activity extracellularly or by transferring to cells that do or do not produce them, i.e., non-cell autonomously. Two separate sequences necessary and sufficient for HP cell exit and entry are in the DNA-binding homeodomain (for review see Di Nardo, 2018). A simple genetic approach cannot therefore be used to study their direct non-cell autonomous activity, as mutation of either sequence alters OTX2 DNA binding and thus also alters cell-autonomous activities. An alternative genetic approach was used to specifically target extracellular OTX2 in order to only abolish non-cell autonomous activity. Conditional mice have been designed to express a neutralizing secreted anti-OTX2 single chain antibody (*scFvOTX2*^*tg/o*^ mice) in a Cre-dependent manner (Bernard 2016). In the retina parvalbumin (PV) is only expressed by RGCs and amacrine cells that do not express *Otx2*. Thus it is anticipated that in *PV*^*Cre*^::*scFvOTX2*^*tg/o*^ mice, OTX2-scFv expressed and secreted from RGCs and amacrine cells will sequester extracellular OTX2 in the vicinity of the producing cells, thus blocking its non-cell autonomous activities. This strategy based on anti-HP scFv *in vivo* secretion has been used with success in several animal models to neutralize extracellular PAX6, ENGRAILED and OTX2 (Lesaffre et al., 2007; Wizenmann et al., 2009; Layalle et al., 2011; Bernard et al., 2016).

We show here that the sequestration of extracellular OTX2 by the OTX2-scFv secreted by RGCs and amacrine cells leads to a significant decrease in visual acuity. This decrease takes place in absence of any observable developmental defects, laminar abnormalities or changes in cell lineages. Electroretinogram (ERG) measurements show normal outer and inner nuclear function, but show a 2-fold increase in amplitude in the response to 20Hz flickers. Taken together, our results provide evidence for a direct non-cell autonomous activity of OTX2 for RGC function.

## Methods and materials

### Production of transgenic mice

*scFvOTX2*^*tg/o*^ and *scFvPAX6*^*tg/o*^ mice were produced by the Institut Clinique de la Souris (Strasbourg, France) as described (Bernard et al., 2016). The mice were crossed with *PV*^*Cre*^ mice obtained from Jackson Labs (stock n°8069).

### Immunoprecipitation and Western blot

Immunoprecipitation and Western blotting were carried out as described (Bernard 2016). Retinas were dissected and suspended in immunoprecipitation lysis buffer (20mM Tris pH8, 120mM NaCl, 1% NP-40, 1mM MgCl2, 5% glycerol, Benzonase nuclease and protease inhibitors). Samples were centrifuged (10min, 20 000*g*) at 4°C and the supernatants incubated overnight at 4°C, with rotation with anti-GFP-coupled magnetic beads (Chromotek). The beads were washed with lysis buffer and 1 M urea before Western blot analysis. Protein extracts were separated on a NuPAGE 4–12% Bis-Tris pre-cast gel (Invitrogen) for 1 h at 200 V and transferred onto a methanol-activated PVDF membrane at 400 mA for 1 h. The membrane was blotted with an anti-MYC antibody (rabbit polyclonal, 1/4000, Sigma-Aldrich C3956) and imaged with a LAS-4000 gel imager (Fujifilm).

### In situ hybridization for OTX2scFV expression

*In situ* hybridization (ISH) was performed on retinal sections using RNAscope technology. Eyes were removed and fixed overnight at 4 °C in buffered 4% paraformaldehyde (PFA) and cryoprotected in 20% sucrose overnight at 4 °C. Forty µm cryostat sections were collected on Superfrost slides, dried and stored at −20 °C. On the day of processing, they were thawed to room temperature for 15 min before performing ISH according to manufacturer’s protocol (Advanced Cell Diagnostics). Briefly, endogenous peroxidase was neutralized with H_2_O_2_, permeabilized with buffered Tween 20 using reagents provided by the manufacturer. Probes designed by Advanced Cell Diagnostics were hybridized for 2 h at 40 °C and the signal amplified in three steps was visualized either with Red (570nm) or Far Red (690nm). The probes *PV* (Mm-PV-C2) and *Myc* (Mm-MYC-C1) were used at 1:50 dilution.

### Visual acuity

The optomotor test of visual acuity was used to screen for possible phenotype differences. Postnatal day (P) 30 mice were used since it was thought that this age would allow sufficient time for scFv expression and secretion. The optomotor test was performed as described in the literature with an optotype of 0.375 c/deg known to elicit robust responses (Torero-Ibad et al., 2011; Bernard et al., 2014). Briefly, mice were placed on an elevated platform centered in a rotating drum and a square wave 100% contrast optotype of 0.375 c/deg was rotated at a speed of 2 rpm. Mice were filmed from above and scored in real time by two observers blind to genotype for head turns in the direction and speed of the moving grating. Any discrepancies between the two observers’ real time score was resolved by analyzing the video. A Mann-Whitney U test was used to compare the genotypes due to unequal variances.

### Retinal histology

Eyes were removed and placed in 4% PFA, 2% ZnCl and 20% isopropyl alcohol (Bernard, 2014). Five µm paraffin sections were cut in the vicinity of the optic nerve and stained for hematoxylin and eosin (Excalibur Pathology, Inc, Oklahoma City, OK). Three sections per retina were analyzed per animal. Total number of cells in the entire GCL (i.e., RGCs and displaced amacrine cells) were counted for each of three sections that included the optic nerve head and averaged. The thickness of the inner nuclear layer (INL) and outer nuclear layer (ONL) proximal to the optic nerve head was measured on the three sections per animal and averaged.

### qPCR for OTX2scFv expression and retinal cell types

RNA was extracted from frozen retina of 30 day-old mice (P30) mice with RNeasy lipid tissue mini kit (Qiagen). cDNA was generated with the Quantitect Reverse transcription kit (Qiagen). Using the 2^-ΔΔCt^ method, sample expression levels were normalized to *hprt* and to *PV*^*Cre*^ mice levels. Primers for the following cell types were used: *Otx2-scFv* (transgene expression), *Brn3A* (ganglion cells), *Chx10* (bipolar cells), *Syntaxin* (amacrine and horizontal cells), *Lim1* (horizontal cells), *Rhodopsin* (rods), *Red Opsin* (L/M cone cells). Since the data are normalized to the value for PVCre mice, the nonparapmetric Mann-Whitney U test was used to evaluate the difference for *Lim1* expression.

### Electroretinogram

Mice were anesthetized with xylazine/ketamine (Imalgene 500 Virbac France, 100mg/kg, Rompun 2% Bayer, 10mg/kg). Tropicamide (Mydriaticum 0.5% Théa, France) was used to dilate the iris and the cornea was locally anesthetized with oxybuprocaine hydrochloride. Gold electrodes were placed in contact with each eye, reference electrodes were placed subcutaneously in the submandibular area, and a ground electrode was placed subcutaneously on the back of the animal. ERG was performed with a mobile apparatus (SIEM Bio-Medical, France) with LED lamps in a Ganzfeld chamber controlled by the VisioSystem software. ERG recordings were obtained from seven *PV*^*Cre*^ and nine *PV-Cre::OTX2-scFv* mice 30-31 days of age. The right and left eye of each mouse was considered to be independent. Mann-Whitney U test was used to evaluate the difference between genotypes for 20Hz flickers.

### Statistical table

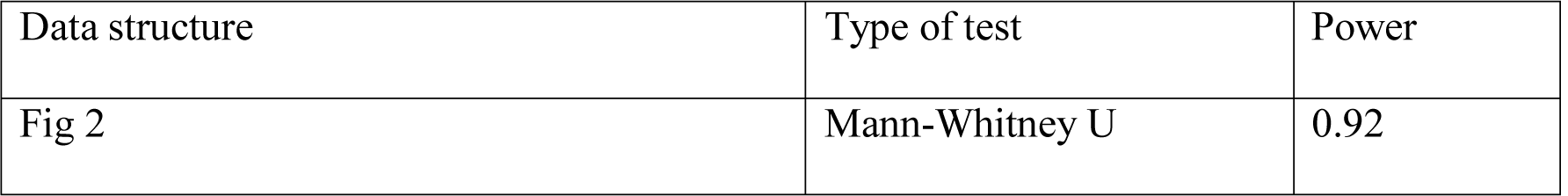
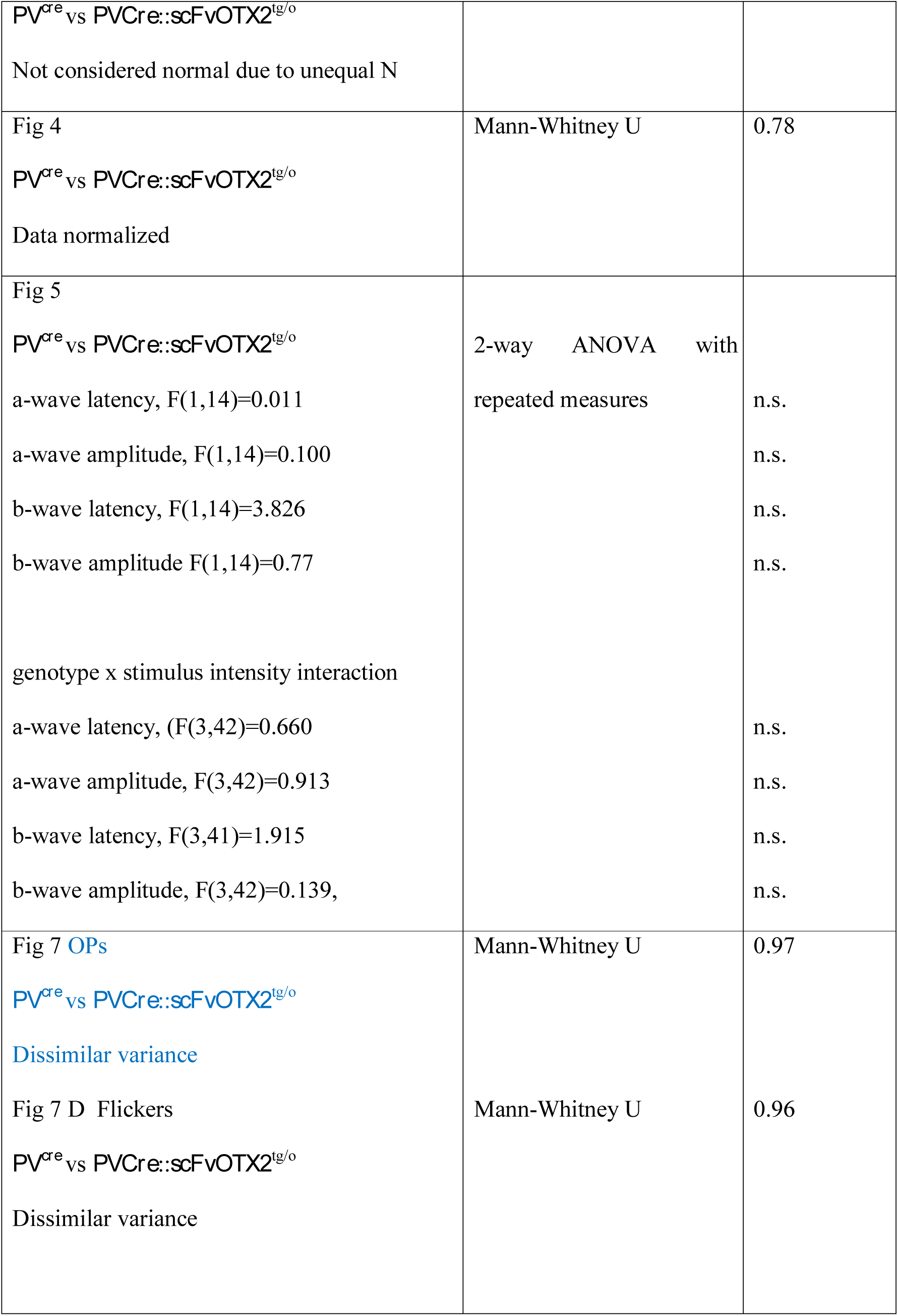

## Results

In the retina, PV is expressed in RGCs and amacrine cells (Haverkamp and Wässle, 2000, Haverkamp et al., 2009). As shown in Fig. 1A, RT-qPCR shows significant expression of the *OTX2-scFv* transgene in the retina of *PV*^*Cre*^::*scFvOTX2*^*tg/o*^ mice and no detectable expression in the retina of *PV*^*Cre*^ mice (raw ct *PVCre* = 38.2±1.7; *PV::Cre*;*scFvOTX2*^*tg/o*^ = 28.9±0.1). Because the antibody is fused with GFP (Bernard et al., 2016), we immunoprecipitated the OTX2-scFv protein from *PV*^*Cre*^::*scFvOTX2*^*tg/o*^ retina using an anti-GFP antibody followed by Western blot analysis. We detected a doublet at the expected 78kD molecular weight, that was not present in extracts from retina of *PV*^*Cre*^ control mice (arrow Fig 1B). *In situ* hybridization (Fig 1C) revealed *PV*-expressing cells in the inner retina in the reported location of PV-containing cells (Haverkamp and Wässle, 2000). In *PV*^*Cre*^::*scFvOTX2*^*tg/o*^ retina each PV-expressing cell also expressed mRNA for the scFv in *PV*-expressing cells as expected (Fig 1D). This demonstrates the expression of the transgene in PV cells and the presence of full-length OTX2-scFv protein in the retina of *PV*^*Cre*^::*scFvOTX2*^*tg/o*^ mice.

**Figure 1.**
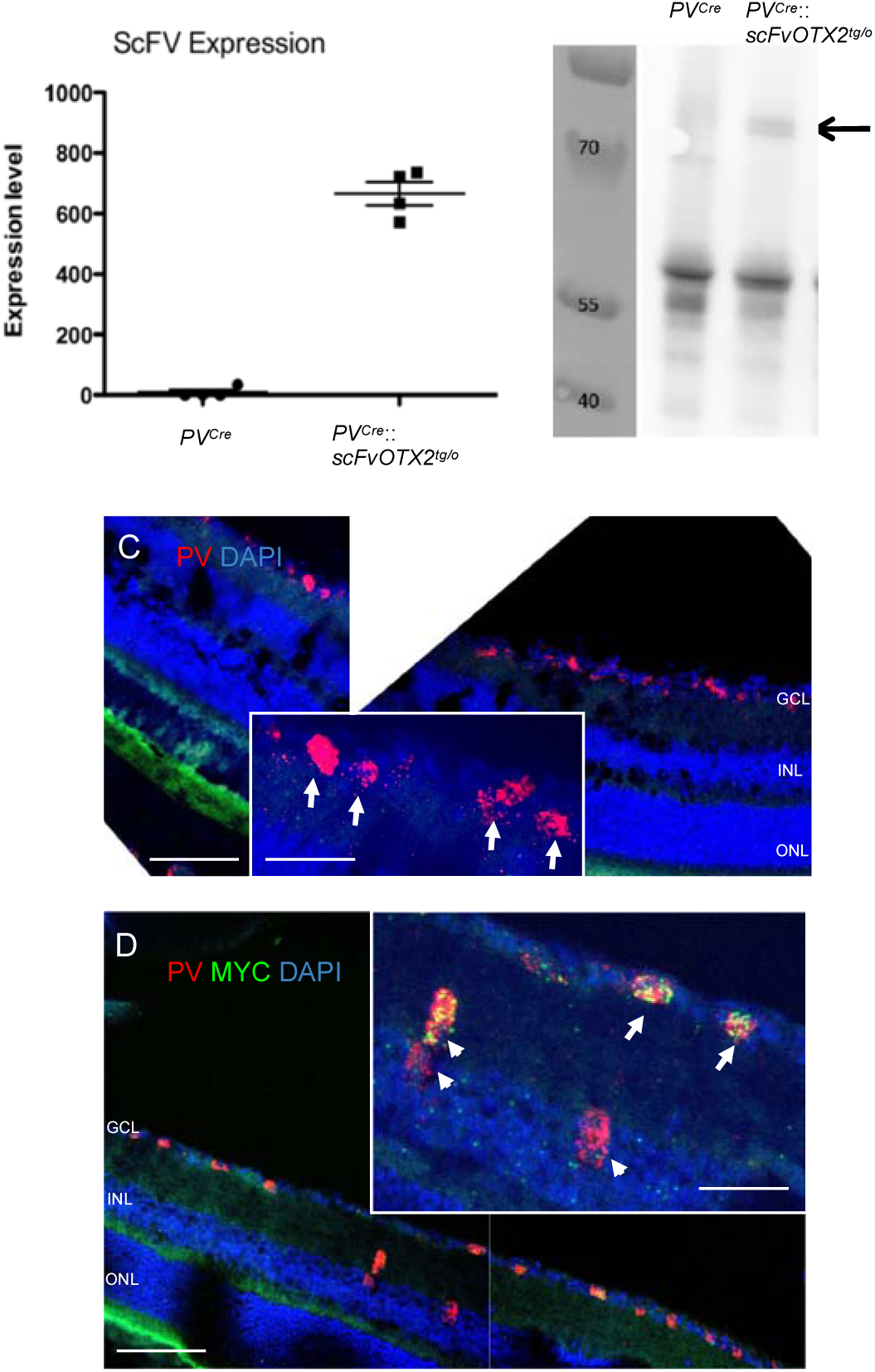
Expression of OTX2-scFV in P30 mouse retinal tissue. A: RTqPCR for OTX2scFV mRNA was carried out on extracts from *PV*^*Cre*^ and *PV*^*Cre*^::*scFvOTX2*^*tg/o*^ mice and raw ct values normalized and converted to arbitrary units (a.u.). B: Retinal lysates from *PV*^*Cre*^ and *PV*^*Cre*^::*scFvOTX2*^*tg/o*^ mice were immunoprecipitated for GFP followed by Western blotting with and anti-MYC antibody. Molecular weight markers are in the left lane in kDa. The middle lane contains the lysate from *PV*^*Cre*^ mice. Right lane contains lysate from *PV*^*Cre*^::*scFvOTX2*^*tg/o*^ mice. The arrow indicates the presence of the bands at the expected migration position of the full-length OTX2scFv-GFP protein. C: *In situ* hybridization using RNAscope Technology of *PV*^*Cre*^ retina shows PV expression (red) in the inner retina. DAPI is in blue. Inset shows details of PV-expressing cells in the GCL. D: *In situ* hybridization using RNAscope Technology of *PV*^*Cre*^::*scFvOTX2*^*tg/o*^ retina. mRNA for PV is red and mRNA for the myc tag of the OTX2scfv is green.. Arrows indicate scFv-expressing PV cells in the GCL i.e., displaced amacrines and/or RGCs; arrowheads indicate scFv-expressing PV cells in the innermost part of the INL in the place of amacrtine cells. Scle bar: 100μm for low magnification; 50μm insets.

One-month-old mice were tested for visual acuity by examining their optokinetic response to a 100% contrast optotype square wave grating of 0.375 c/deg spatial frequency. Bernard et al (2014) reported that this spatial frequency was most robust in revealing differences in mice hypomorph for OTX2 activity and the number of head movements in response to this optotype is sensitive to changes in visual acuity due to loss of RGCs (Torero Ibad et al., 2011). A significant reduction in visual performance in the *PV*^*Cre*^::*scFvOTX2*^*tg/o*^ mice was observed compared to *PV*^*Cre*^ littermates (Fig 2). As an additional control, we also evaluated *PV*^*Cre*^::*scFvPAX6*^*tg/o*^ mice. PAX6 homeoprotein is reported to also have non-cell autonomous activities (Di Lullo et al., 2011; Kaddour et al., 2019), and these mice showed no visual acuity loss. These results demonstrate that neutralizing extracellular OTX2 in the retina leads to decreased visual acuity at P30 but the present results do not allow us to determine if this persists at later ages.

**Figure 2.**
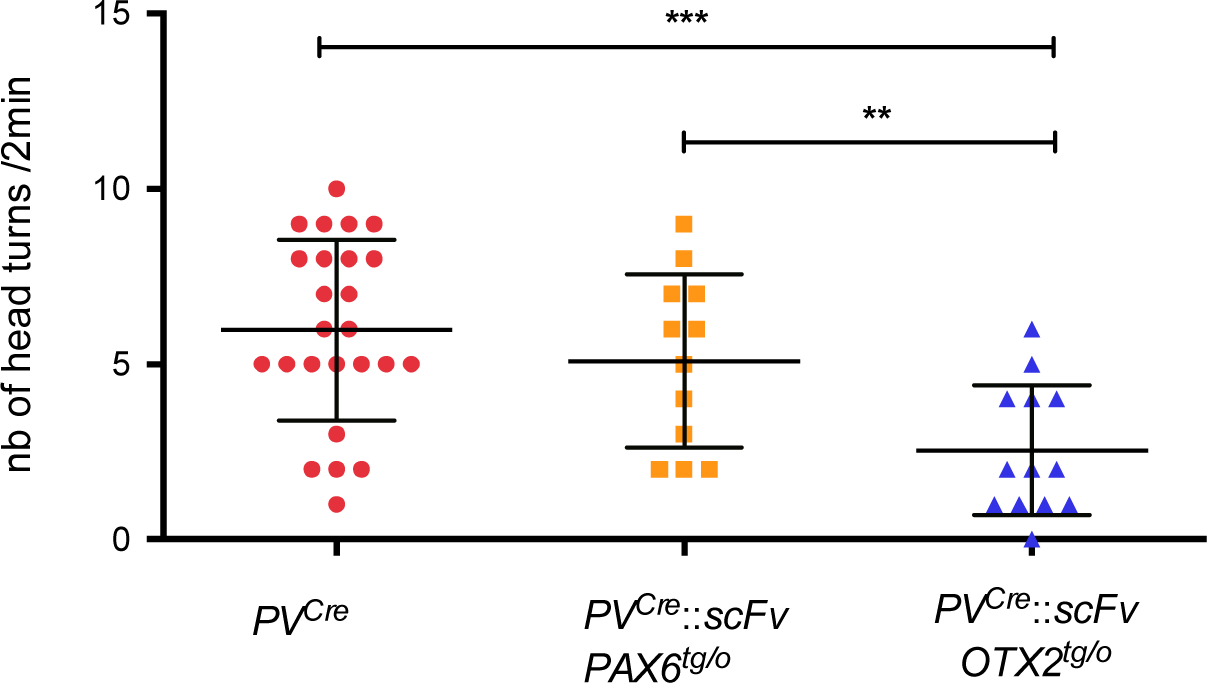
Optomotor test monitoring visual acuity. Thirty-day-old mice were subjected to the optomotor test using an 100% contrast optotype of 0.375c/deg. *PV*^*Cre*^ and *PV*^*Cre*^::*scFvPax6*^*tg/o*^ mice made an average of about five head turns during the 2 min test period. *PV*^*Cre*^::*scFvOTX2*^*tg/o*^ mice made significantly fewer head turns revealing reduced visual acuity. **p<0.01 or ***p<0.001, Mann-Whitney U test, two tailed. N=25 for *PV*^*Cre*^, N=12 for *PV*^*Cre*^::*scFvPax6*^*tg / o*^ and N=3 for *PV*^*Cre*^::*scFvOTX2*^*tg/o*^

In search of a cellular correlate, we analyzed retinal structure by hematoxilin and eosin staining of sections from 1month-old mice. There was no difference in thickness of the outer nuclear layer (ONL, photoreceptors) nor of the inner nuclear layer (INL; horizontal, bipolar, amacrines and Müller glial cells) between *PV*^*Cre*^::*scFvOTX2*^*tg/o*^ and *PV*^*Cre*^ littermates (Fig 3). Nor could we find a difference in the number of cells in the ganglion cell layer (GCL) containing RGCs and displaced amacrine cells. These results thus show no gross structural abnormalities in the retina of *PV*^*Cre*^::*scFvOTX2*^*tg/o*^ mice nor obvious differences in the number of cells in the ONL, INL or GCL.

**Figure 3.**
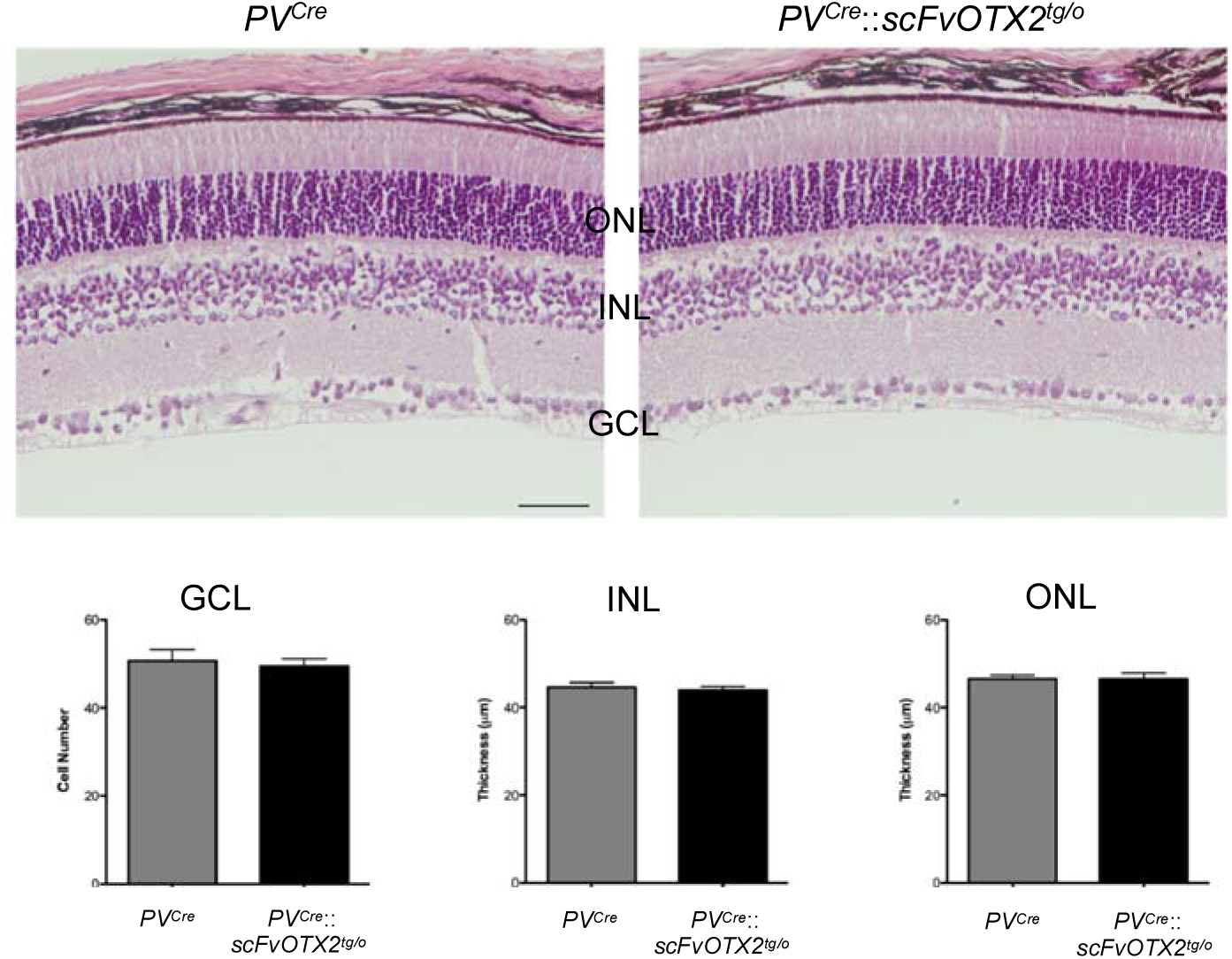
Expression of OTX2scFv does not alter retinal organization in the P30 mouse. Upper: all retinal layers organization and approximate sizes are similar in retina from mice expressing OTX2scFv and their *PV*^*Cre*^ littermates. Lower: the number of cells in the GCL, the INL thickness and the ONL thickness are similar between the two genotypes. Scale bar: 50μM. Abbreviations: GCL, ganglion cell layer; INL, inner nuclear layer; ONL, outer nuclear layer. N=8 for *PV*^*Cre*^ and N=6 for *PV*^*Cre*^::*scFvOTX2*^*tg/o*^ mice.

Quantitative RT-PCR was reported to faithfully reflect retinal cell number (Torero-Ibad et al., 2011). We observed no differences between P30 *PV*^*Cre*^::*scFvOTX2*^*tg/o*^ and *PV*^*Cre*^ littermates for expression of mRNAs for *Brn3A* (RGCs), *syntaxin* (amacrine and horizontal cells), *Chx10* (bipolar cells), *rhodopsin* (rod photoreceptors) and *red/green opsin* (long/medium wavelength cone photoreceptors) (Fig. 4). A small increase in *Lim1* expression by horizontal cells was observed which might reflect a non-cell autonomous repressor activity of OTX2 on *Lim1* expression (Nakano et al., 2000; Puelles et al., 2006). Thus, based on these markers, histological studies and cell counting, it can be concluded that all cell types involved in retinal visual processing are present in normal numbers.

**Figure 4.**
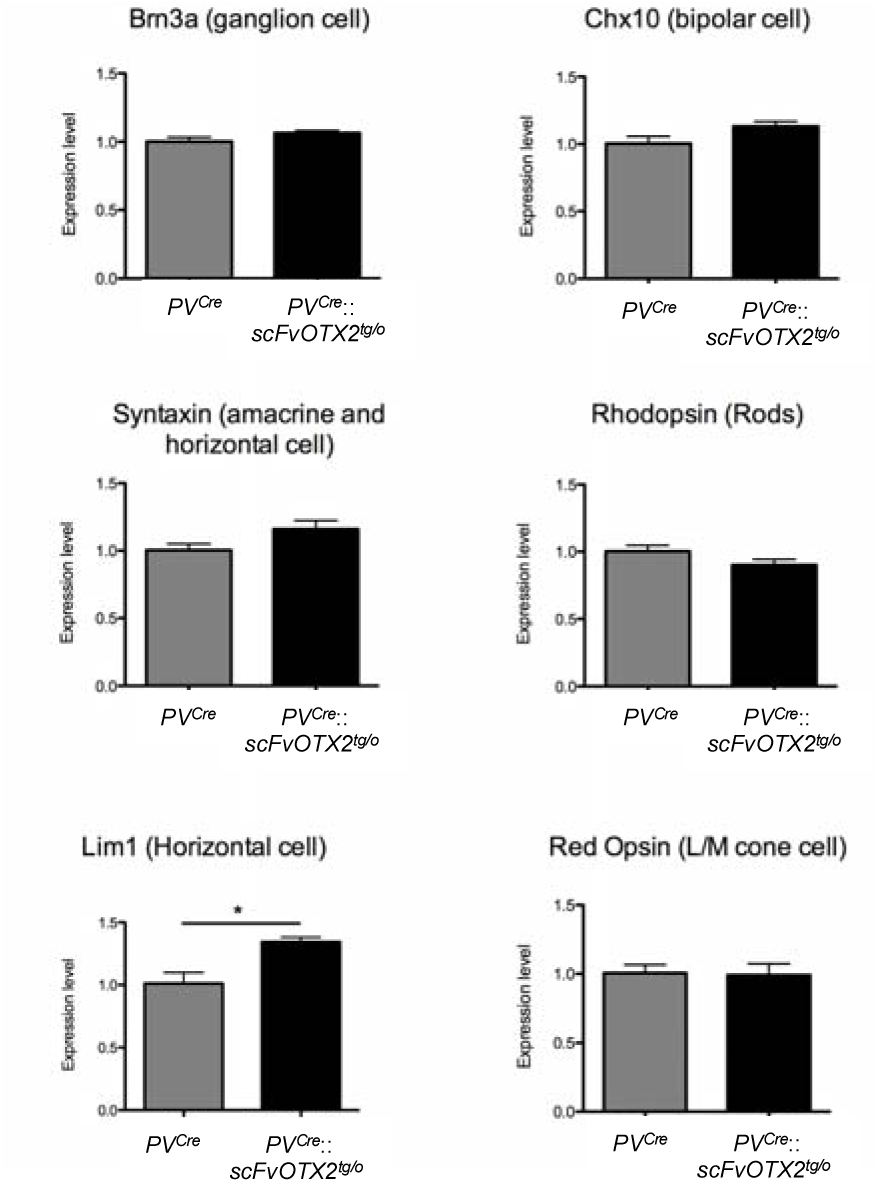
Expression of retinal cell type-specific genes in P30 mice of the two genotypes. Genes specific of RGCs, bipolar cells, amacrine cells, rods and cones show similar expression levels in the two genotypes. *Lim1* mRNA expression in horizontal cells is significantly increased in the OTX2-scFv expressing mice. P<0.05, Mann-Whitney U-test, two -tailed. N=4 for each genotype.

To determine if there was a functional defect in photoreceptors and/or bipolar cells we turned to ERG. The negative a-wave reflects photoreceptor hyperpolarization in response to light. The amplitude is related to photoreceptor number and the implicit time to photoreceptor physiology (Lamb and Pugh, 2004; Hood and Birch, 1992; see Weymouth and Vingrys, 2007 for review). Figure 5A,B shows representative scotopic traces for retinas from *PV*^*Cre*^::*scFvOTX2*^*tg/o*^ and *PV*^*Cre*^ mice. A- and b-wave implicit times and amplitudes were plotted against log intensity (Fig 5C-F). Two-way ANOVA with repeated measures revealed no significant differences between the two genotypes (a-wave latency, F(1,14)=0.011; a-wave amplitude, F(1,14)=0.100; b-wave latency, F(1,14)=3.826; b-wave amplitude F(1,14)=0.77, all nonsignificant) nor for genotype x stimulus intensity interaction (a-wave latency, (F(3,42)=0.660; a-wave amplitude, F(3,42)=0.913; b-wave latency, F(3,41)=1.915; b-wave amplitude, F(3,42)=0.139, all nonsignificant). The absence of a difference in a-wave amplitude or implicit time in the *PV*^*Cre*^::*scFvOTX2*^*tg/o*^ mice compared to *PV*^*Cre*^ littermates indicates normal number and function of rod photoreceptors (Fig. 5 and 7). The b-wave is mainly driven by bipolar activity and the amplitude reflects number and the implicit time bipolar function. Both were indistinguishable between *PV*^*Cre*^::*scFvOTX2*^*tg/o*^ mice and *PV*^*Cre*^ littermates indicating normal number and response of bipolar cells. The b-wave laency and amplitude under photopic conditions was similar in *PV*^*Cre*^::*scFvOTX2*^*tg/o*^ mice and *PV*^*Cre*^ littermates (Fig 6). Interestingly, we observed a significant decrease in the amplitude in oscillatory potential 4 (OP4) in *PV*^*Cre*^::*scFvOTX2*^*tg/o*^ mice (Fig 7A,B). There were no changes in earlier or later OPs. Additionally, for the flicker response, the *PV*^*Cre*^::*scFvOTX2*^*tg/o*^ mice had an increase of about 33% in amplitude at 10 Hz and at 20Hz the response significantly increased by 2-fold (Fig 7C,D). Both of these results suggest inner retinal dysfunction at the level of amacrine cells and/or RGCs.

**Figure 5.**
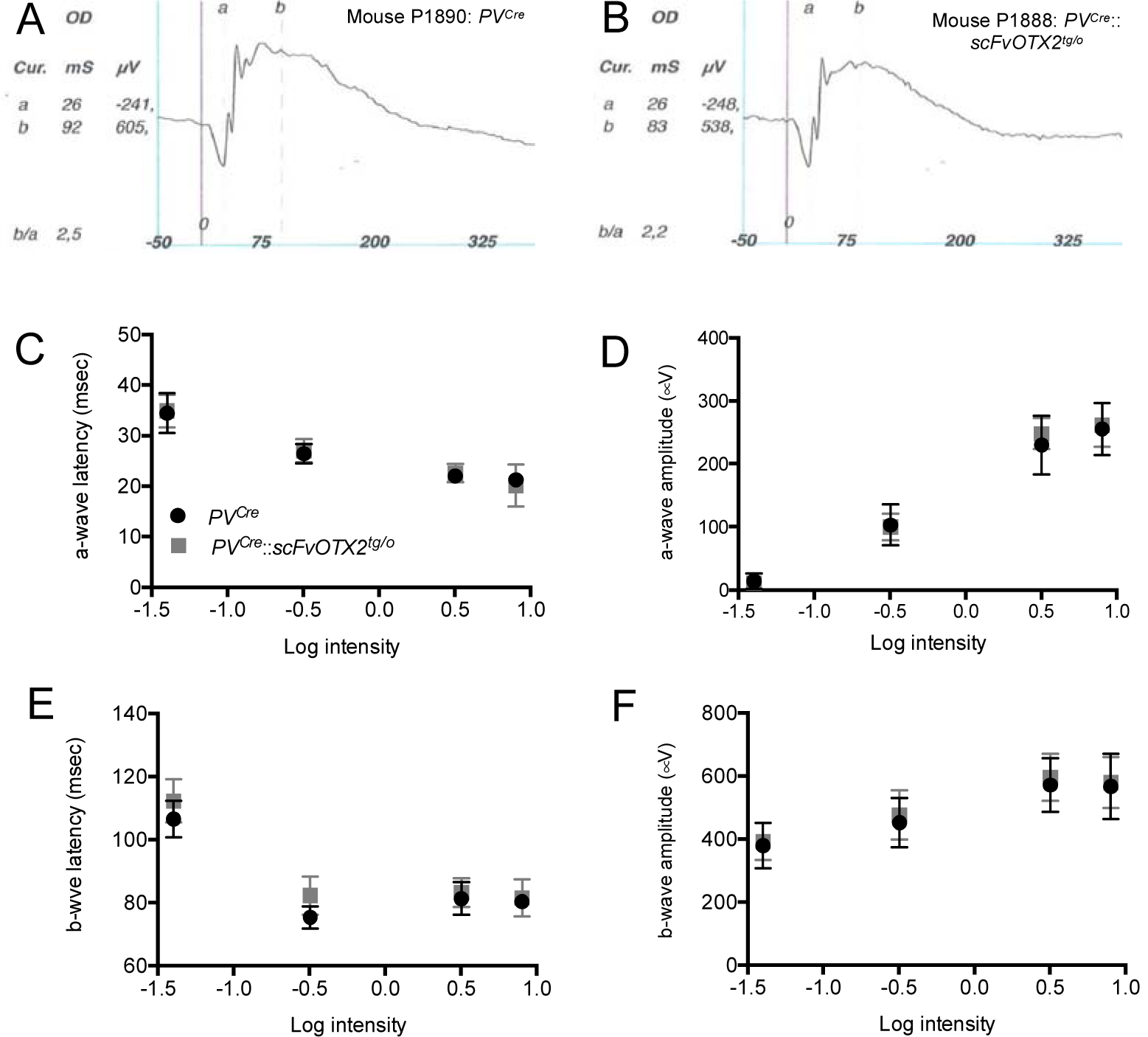
ERG P30 mice expressing OTX2scFV and their PV^Cre^ littermates. A,B: Rrepresentative traces under scotopic conditions of the right eye at 3.19cds/m^2^. 20Hz flickers of the left eye. C-F: Scotopic ERG parameters plotted against log intensity. Two-way ANOVA for repeated measures showed no significant differences based on genotype nor genotype by intensity interaction (see text).

**Figure 6.**
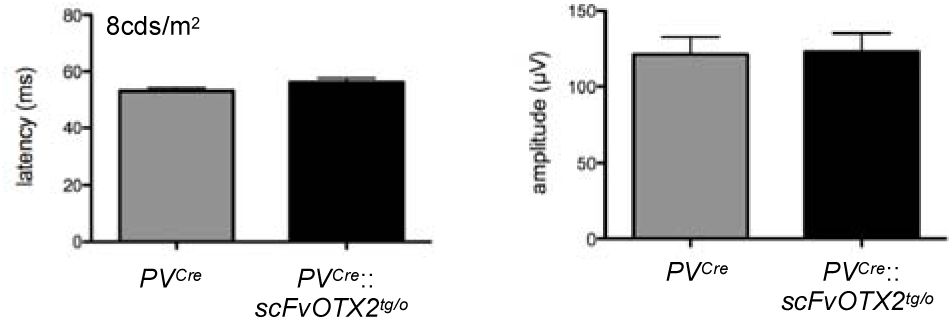
B-wave amplitude in photopic lighting in the two genotypes of one-month-old mice. N=7 for *PV*^*Cre*^ and N=9 for *PV*^*Cre*^::*scFvOTX2*^*tg/o*^ mice.

**Figure 7.**
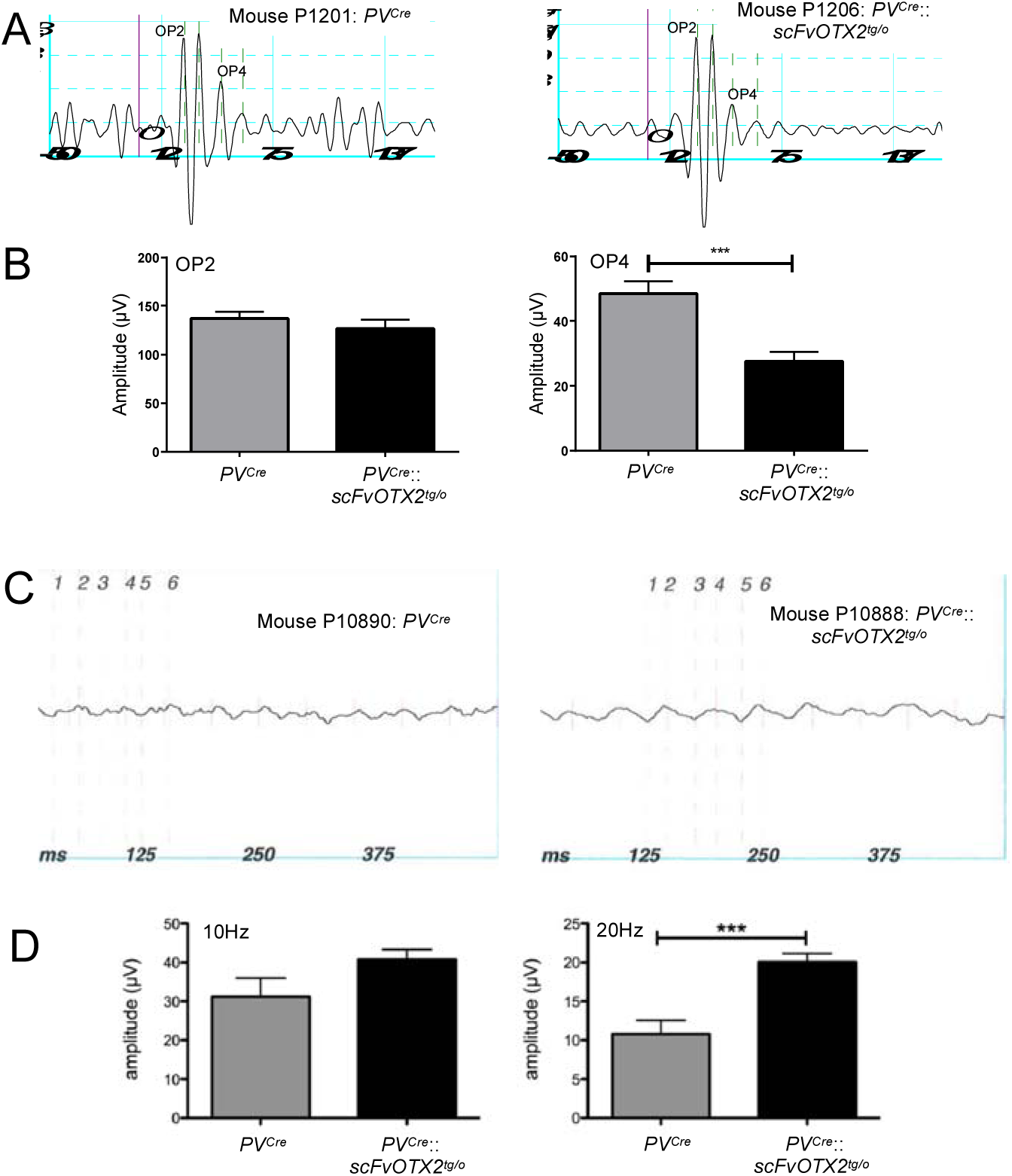
Inner retinal function. A: Extracted oscillatory potential traces of a *PV*^*Cre*^ (left) and *PV*^*Cre*^::*scFvOTX2*^*tg/o*^ (right) mouse. B: The amplitude of an early OP (OP2) is not altered in the *PV*^*Cre*^::*scFvOTX2*^*tg/o*^ mice, while the amplitude of a late OP (OP4) is significantly reduced by about 50%. C: Represcentative traces of 20Hz flickers of the left eye of mice of the two genotypes. D: The amplitude of the flicker response to 10Hz stimulation after light adaptation is slightly increased and at 20Hz the response is significantly doubled in amplitude. Lower: there was no difference in the implicit time of the flicker responses between mice expressing OTX2scFv and their *PV*^*Cre*^ littermates. P<0.005, Mann-Whitney U-test, two-tailed. N=7 for *PV*^*Cre*^ and N=9 for *PV*^*Cre*^::*scFvOTX2*^*tg/o*^ mice.

## Discussion

OTX2 homeoprotein is part of the HP family of transcription factors that transfer between cells and have neuroprotective effects in the brain (Sugiyama et al., 2008; Torero-Ibad et al., 2011; Rekaik et al., 2015; see DiNardo et al, 2018 for review). We were interested in assessing the importance of the direct non-cell autonomous activity of OTX2 in the retina since the Otx2 locus is silent in cells of the GCL, yet they contain OTX2 protein (Rath et al., 2007; Sugiyama et al., 2008). Since the amino acid sequences for OTX2 to exit and enter cells lie in the DNA binding homeodomain, mutations in these sequences might alter DNA binding and thus cell autonomous activity. Bernard et al., (2016) used a genetic approach to neutralize extracellular OTX2 by expressing a secreted anti-OTX2 single chain antibody. Here, we used this approach to drive expression of the OTX2-scFv in cells which do not themselves produce OTX2, thus eliminating a putative cell autonomous effect of the antibody. PV is reported to be in AII amacrines in rats, cats, bats and rabbits (Lee et al., 2004). In the mouse, direct immunofluorescence showed low PV expression in amacrines and displaced amacrines and strong PV expression in RGCs (Haverkamp and Wässle, 2000). A more sensitive transgenic approach using the PV promoter to drive expression of enhanced GFP (Haverkamp et al., 2009) confirmed this expression pattern and showed that PV was expressed in some amacrines in the INL with notably strong expression in RGCs (Haverkamp et al., 2009). Kim and Jeon (2006) and later Yi et al., (2012) using single cell iontophoresis and retrograde tracing followed by PV localization showed that at least eight different RGC types express PV. Thus, we anticipated widespread expression of the scFv throughout the GCL and inner INL. It is interesting to note that the ISH results showed scFv expression in fewer cells than expected, but it was sufficient to neurtralize extracellular OTX2 and cause functional and vision changes.

Parvalbumin starts to be expressed in RGCs around P11, after the birth of all retinal cell types (Cepko et al., 1996; Young, 1985). Thus, it is expected that interference with OTX2 non-cell autonomous activity starting at the earliest at P11 would have no effect in establishing the normal layers and structure of the retina, as we observe. It was reported that in *Otx2* hypomorph mice, in which both cell autonomous and non-cell autonomous phenotypes can be expected, there were fewer photoreceptors at P30, but those that were present have normal a-wave implicit time (Bernard et al., 2014). In *PV*^*Cre*^::*scFvOTX2*^*tg/o*^ mice, the presence of normal numbers of OTX2-expressing photoreceptors and bipolar cells and their normal ERG a-wave and b-wave activity, respectively, confirms that OTX2 cell autonomous activity in these cells is not affected by expression of the secreted OTX2-scFv. It further suggests that OTX2 non-cell autonomous activity on these cells does not contribute measurably to their survival and function, at least at one month of age.

It is notable that we observed no changes in retinal structure or organization in the *PV*^*Cre*^::*scFvOTX2*^*tg/o*^ mouse retina and that both ONL and INL function at least in terms of ERG a- and b-wave activity was comparable to *PV*^*Cre*^ littermates and yet the *PV*^*Cre*^::*scFvOTX2*^*tg/o*^ mice had a deficit in visual acuity. How might this occur? This was not simply due to the expression of an scFv since *PV*^*Cre*^::*scFvPAX6*^*tg/o*^ mice did not show any loss of visual acuity compared to *PV*^*Cre*^ mice. PV is expressed in other regions of nervous system, and could interfere with OTX2 non-cell autonomous activity in these regions, thus contributing to the reduction of visual acuity. The optokinetic response does not depend on visual cortex since large lesions of posterior cortex do not diminish visual acuity, indicating that the brain circuits for this response are entirely subcortical (Douglas et al., 2005). The basic circuit for the optokinetic response is retinal projection to the nucleus of the optic tract that sends input to the reticular tegmental nucleus of the pons with output going to the vestibular nucleus driving the oculomotor nucleus (Wada et al., 2014). None of these structures are known to express OTX2 and only the vestibular nucleus has PV cells. Thus, the only structure in the basic circuit that has both PV and OTX2 expression is the retina.

We propose that the loss of non-cell autonomous signaling OTX2 induces inner retinal physiological dysfunction as suggested by the altered OP and flickers response. OPs are an indication of inner retinal function and appear to be driven mainly by rod activity (Wachtmeister, 1998; Lei et al., 2006). OP ERG has been used to evaluate inner retina function in both human and and experimental glaucoma (Rangaswamy et al., 2006; Gur et al., 1987). Early OPs have been attributed to GABAergic inhibitory activity in the ON-pathway while later OPs have been pharmacologically characterized as being generated by glycinergic inhibitory amacrine feed-back synapses in the OFF-pathway (see Wachtmeister, 1998). Flickers reflect cone pathway activity (Tanimoto et al., 2015) and the cone photoreceptor function and cone bipolar function as measured in the photopic a- and b-wave ERG is not compromised in the *PV*^*Cre*^::*scFvOTX2*^*tg/o*^ retinas. Thus, the altered flicker responses that we observe represents a change in cone pathway activity at the level of the inner retina. Retinal Ganglion Cell dysfunction and degeneration in a model of glaucoma greatly alters the flicker response (Grozdanic et al., 2003) and the flicker ERG has been used to assess RGC function and hemodynamic changes in the mouse retina (Chou et al., 2019). The flicker response in macaques has been shown to be largely driven by spiking inner retinal neurons since it is increased at some frequencies by TTX and NMDA (Viswanathan et al., 2002). Here, we observed the same direction of change in *PV*^*Cre*^::*scFvOTX2*^*tg/o*^ mice at 10Hz and significantly at 20Hz. OPs and the flickers are elicited by different stimuli. The OPs are part of the b-wave response elicited in our case by scotopic flash. The flickers are elicited by flicker stimuli in light adapted conditions. Thus, two independent stimuli in different light adaptation conditions reveal inner retinal dysfunction when there is a decrease in OTX2 signaling in the retina.

We propose that non-cell autonomous OTX2 signaling in the inner retina is affected without loss of RGCs or amacrine cells in the GCL in the *PV*^*Cre*^::*scFvOTX2*^*tg/o*^ mouse retina. What could be the nature of this signaling? Homeoproteins including OTX2 are capable of transferring between cells and, in addition to their transcriptional activity, can stimulate protein translation, alter chromatin remodeling and repress transposable element expression (Blaudin de Thé, 2018 and Di Nardo et al., 2018 for review). OTX2 from the retina can reach inhibitory PV cells in V1 cortex and interference with its non-cell autonomous activity perturbs the normal opening and closing of the critical period for ocular dominance plasticity by altering the balance of excitation and inhibition (Sugiyama et al., 2008; Bernard et al., 2016). The change in flicker amplitude we recorded in the *PV*^*Cre*^::*scFvOTX2*^*tg/o*^ mouse retina may be related to a change in the excitation/inhibition of the inner retinal circuitry. It was reported that extracellular OTX2 neutralization in the retina can retard the opening of critical period plasticity in V1 cortex (Sugiyama et al. 2008). Since visual acuity measured by the optomotor test is independent of the visual cortex, this means that OTX2 signaling in the retina has separate physiological consequences both on subcortical and cortical visual circuits.

Non-cell autonomous signaling by the homeoproteins ENGRAILED 1/2 is important for RGC axon guidance in Xenopus and chick tectum and this signaling involves stimulating ATP production and release from the RGC growth cone and subsequent activation of adenosine A1 receptor (Brunet et al., 2005; Wizenmann et al., 2009; Stettler et al., 2012). Retinal Ganglion Cells have particularly high energy demands and energy homeostasis is critical for their function, and energy deficits are hypothesized to underlie their dysfunction and degeneration in diabetes, glaucoma and optic neuropathies (Osborn et al., 2013; Yu et al., 2013; Ito and Di Polo, 2017). Bipolar cell degeneration in *Otx2* hypomorph mice has been attributed to mitochondrial dysfunction that can be reversed by exogenous OTX2 (Bernard et al., 2014; Kim et al., 2015). While our finding could possibly be explained by a metabolic defect, it remains to be determined if RGC mitochondrial energetics are altered in the *PV*^*Cre*^::*scFvOTX2*^*tg/o*^ retina and whether this affects inner retina activity. Because displaced amacrine cells and RGCs in the GCL, major components of the inner retinal circuit, do not themselves express the *OTX2* gene but are capable of taking up exogenous OTX2 (Sugiyama et al., 2008; Torero-Ibad et al., 2011), it can be proposed that. interfering *in vivo* with OTX2 non-cell autonomous activity at the level of the mature retina leads to an alteration in inner retinal cell functions in early life and causes the observed deficit in visual acuity. Future studies will be required to establish if these functional changes a longer lasting.

## Acknowledgements

We thank Ms. Mélanie Leboeuf and Drs. Anne-Cécile Boulay and Martine Cohen-Salmon for assistance with the RNAscope experiments.

